# Assessing the impact of boldine on the gastrocnemius using multiomic profiling at 7 and 28 days post-complete spinal cord injury in young male mice

**DOI:** 10.1101/2022.08.17.503230

**Authors:** Luke A. Potter, Carlos A. Toro, Lauren Harlow, Kaleen M. Lavin, Christopher P. Cardozo, Adam R. Wende, Zachary A. Graham

## Abstract

Spinal cord injury (SCI) results in rapid muscle loss. The mechanisms of muscle atrophy have been well-described but there is limited information specific to SCI. Exogenous molecular interventions to slow muscle atrophy in severe-to-complete SCI have been relatively ineffective and the wide-ranging physiologic response to SCI requires the search for novel therapeutic targets. Connexin hemichannels (CxHC) allow non-selective passage of small molecules into and out of the cell. Boldine, a CxHC-inhibiting aporphine found in the boldo tree (*Peumus boldus*), has shown promising pre-clinical results in slowing atrophy during sepsis and dysferlinopathy. We administered 50 mg/kg/d of boldine to spinal cord transected mice beginning 3 d post-injury. Tissue was collected 7 and 28 d post-SCI and the gastrocnemius was used for multiomic profiling. Boldine did not prevent body or muscle mass loss but attenuated SCI-induced changes in the abundance of proline, phenylalanine, leucine and isoleucine, as well as glucose, 7 d post-SCI. SCI resulted in the differential expression of ~7,700 and ~2,000 genes at 7 and 28 d, respectively, compared to sham animals, with enrichment for pathways associated with ribosome biogenesis, translation and oxidative phosphorylation. Boldine altered the expression of ~150 genes at 7 d and ~110 genes at 28 d post-SCI. Methylation analyses highlighted distinct patterns at both 7 and 28 d following SCI both with and without boldine. Taken together, boldine is not an efficacious therapy to preserve body and muscle mass after complete SCI, though it preserved or attenuated SCI-induced changes across the metabolome, transcriptome and methylome.

## Introduction

Muscle atrophy is a highly-coordinated process that occurs during periods of disuse. Sustained unloading elevates the rate of protein degradation (1), disrupts protein synthesis by reducing RNA translation efficiency (2) and promotes dietary anabolic resistance (3). These factors can all occur in parallel with the resulting outcomes of reduced size, function and health of skeletal muscle. Reductions in muscle mass have metabolic implications, as it is the principal site of insulin-mediated glucose clearance (4), and functional implications, as disuse reduces strength and power (5). One of the most severe forms of muscle atrophy occurs after spinal cord injury (SCI). Severe SCI such as ASIA Impairment Score (AIS) A/B results in long-standing paresis/paralysis in the muscles below the area of spinal injury with rapid loss in size of the large muscle groups of the hips and legs, losses in muscle mass of the trunk, and for those with higher (e.g., cervical) injuries, chest and arm muscles. Additionally, severe injuries result in less functional recovery over time when compared to less severe AIS C/D injuries (6, 7). Muscle atrophy post-SCI is rapid; our group has consistently observed losses in mass observed by 7 d in pre-clinical models of complete SCI with slower but additional losses seen out to 28-84 d post-injury (8–11).

Muscle mass can be maintained or increased after SCI using interventions that load muscle, such as neuromuscular electrical stimulation/functional electrical stimulation [NMES/FES; (12–15)] or body-weight supported treadmill training (16). Molecular interventions have shown little efficacy in preserving muscle mass to sham control levels in pre-clinical models of severe and complete SCI (17) even when muscle is exposed to potent stimuli such as FES, testosterone, or the combination of the two (18). However, these interventions have led to 7-15% increases in muscle mass compared to SCI-only controls, though the response varies based on the muscle (18–21). A potent myostatin inhibitor was not able to preserve hindlimb muscle mass 56 d after a complete SCI despite gains in muscle mass above the area of lesion (8). Similar outcomes have been noted for other anabolic agents such as testosterone and nandrolone (19, 22). Reasons for these surprising results are unknown but presumably include activation of multiple pathways for muscle atrophy after SCI such that no single drug can block the net effect. The mechanisms that drive the initial loss of muscle mass during acute disuse have been well-described (23–25) but there remain some intriguing targets that have not been extensively studied. Connexin hemichannels (CxHC) are a potential target known to drive muscle atrophy (26–31).

CxHC are non-selective pore proteins composed of hexameric connexin proteins localized, generally, to cell membranes of many different cell types. When open in the hemichannel form they allow efflux of small molecules such as glutamate, ATP and potassium out of the cell, while also allowing influx of ions such as Ca^2+^ and Na^+^ from the extracellular space (32, 33). CxHC can bind with other CxHC from adjacent cells to form gap junctions (GJ). GJ and CxHC are essential in developing muscle for cellular communication during differentiation, regeneration and development (26). As muscle differentiation proceeds and muscle fibers begin to be innervated, the connexins expressed in developing muscle, Cx43 and 45, are downregulated and are not expressed at the sarcolemma of healthy adult skeletal muscle fibers (26, 34). When exposed to insults that may lead to muscle atrophy, such as a sciatic nerve transection (27), dexamethasone administration (28) and endotoxemia/sepsis (35), CxHCs reappear on the sarcolemma of fast-twitch muscle fibers. Appearance of sarcolemmal CxHC lowers membrane potential (30) and activates the inflammasome (27). Interestingly, disuse from hindlimb unloading does not result in increased CxHC appearance (30), demonstrating additional mechanisms outside of disuse drive CxHC cellular translocation. Conditionally knocking out Cx43 and 45 in skeletal muscle can reduce muscle atrophy following denervation by a sciatic nerve transection (27), and inhibiting their activity with molecular interventions has positive effects on muscle health (29, 30, 35, 36). A promising molecule is boldine, an alkaloid derived from the Chilean boldo tree (*Peumus boldus*). Boldine can inhibit CxHC activity without blocking GJ function (37), and has been shown to prevent sepsis-induced increases in cytosolic calcium concentrations and cytokine-induced small molecule dye uptake in isolated muscle fibers (35). Furthermore, 8 wks of boldine treatment in mice with dysferlinopathy improved motor function, reduced skeletal muscle fat accumulation and rescued aberrant dysferlin protein expression (29).

There is little information regarding how SCI affects large scale molecular profiles in mouse skeletal muscle. Additionally, it is not well understood if CxHC have a role in paralyzed skeletal muscle after SCI. It has been shown that a complete thoracic spinal cord transection leads to increased sarcolemmal expression of Cx43 at 56 d after injury in the gastrocnemius of young adult rats (27). However, it is unknown whether inhibition of CxHC activity in the acute and subacute timeframe protects muscle mass or alters the molecular profiles of muscle after SCI. Therefore, in this study, we aimed to: 1) describe how SCI affects the metabolome, transcriptome and methylome of adult mouse skeletal muscle; 2) determine if boldine could prevent losses in body and muscle mass following a complete thoracic SCI; and 3) determine if boldine could alter molecular profiles in a way that could improve muscle health and/or function.

## Methods

### Animals

Male C57BL/6NCrl mice aged 4 months were used for these studies. Mice were purchased from Charles River and kept in an AAALAC-accredited animal facility at the James J. Peters VAMC for a minimum of 2-wks before surgery. Animals were kept on a 12:12 light-dark cycle with ad libitum access to standard chow (LabDiet rodent Diet #5001) and water. Animals were randomly selected for a 7 or 28 d post-surgery timepoint, then further randomized for a laminectomy (Sham) or laminectomy + complete T10 spinal cord transection with vehicle (SCIv) or boldine (SCIb) treatment. 32 animals were used for this study, with n=4 for both 7 and 28 d Sham groups and n=6 for all 7 and 28 d SCI groups. All studies were reviewed and approved by the IACUC of the James J. Peters VAMC (protocol #: CAR-16-54).

### Laminectomy and spinal cord transection

Detailed methods for the laminectomy and spinal cord transection used by our group have been previously published (8). In brief, mice were weighed and then placed under deep anesthesia using inhalation of 2-3% continuous-flow isoflurane. Hair along the back of the animal was shaved and the skin cleaned with 70% ethanol and a 2% betadine solution. A sterile scalpel was then used to make an incision from T7-11 and the spinal column was exposed by blunt dissection and removal of the para-vertebral muscles. The vertebral arch of the T10 vertebral body was removed by cutting the lateral processes with sharp surgical scissors and gently lifting with fine forceps. With the dura exposed, the incision site for the sham animals was promptly closed in layers using sutures for the muscle layer and surgical staples for skin. For the SCI animals, a micro-scalpel was passed through the spinal cord. The transection site was inspected for residual tissue bridges, which were cut by a second pass of the scalpel when present. An inert gel foam was placed in between the ends of the severed spinal cord and the incision site was sutured closed as above, followed by placing the animals in a clean cage. Clean cages included Alpha-dri+ bedding, standard chow placed on the floor for paralyzed animals, peanut butter and Bio-Serv fruit crunch treats. All animals were single-housed for the remainder of the study.

### Post-operative care and boldine administration

Animals were kept on warming pads heated to 37°C with recirculating water for 24 h post-surgery. They were given a daily injection of ketophen (5 mg/kg) and Baytril (5 mg/kg) for 3 d post-surgery and a total daily volume of 1 ml of lactated Ringer’s solution for 7 d to prevent dehydration. Bladders were expressed 2 times per day, and animals were carefully monitored to ensure no signs of an incomplete transection (hindlimb movement, spasms, grooming, manifestations of pain, etc.). Boldine was administered in a 1.0 g bolus of peanut butter at a dose of 50 mg/kg/d starting at 3 d post-injury. Animals were familiarized with peanut butter for a week prior to surgery. All animals consumed 100% of their daily peanut butter by day 3 post-surgery and continued to do so throughout the remainder of the study.

### Body and tissue mass

Mice were euthanized at 7 or 28 d post-surgery. Body mass was recorded before mice were placed under deep anesthesia using inhalation of 2-3% continuous-flow isoflurane. Muscles [soleus, plantaris, gastrocnemius, tibialis anterior (TA) and extensor digitorum longus (EDL), biceps, triceps] were carefully excised, wet-weighed, then flash frozen in liquid nitrogen-cooled isopentane. Animals were sacrificed using a combination of blood collection via ventricular puncture and cervical dislocation while under deep anesthesia. The gastrocnemius was used as it is large enough after severe/complete SCI (~60 mg for a 25 g mouse) for one single muscle to be used for all downstream multiomics platforms.

### Metabolomics and analysis

~10 mg of the left gastrocnemius containing equal portions of medial and lateral heads were sent on dry ice to West Coast Metabolomics, a regional NIH Resource Core located at the University of California-Davis, for untargeted primary metabolomics analyses using GC-TOF mass spectroscopy. Complete sample processing, data acquisition and data processing have been reported in detail (38). Our group has used these parameters to interrogate changes in muscle from female mice at 7 and 28 d post-SCI (10) and others have used these parameters to describe other metabolic changes in mouse skeletal muscle (39). We analyzed peak spectra for each metabolite following normalization to the median, log transformation and pareto scaling. All spectra were matched to known metabolites using the BinBase algorithm (40) while unconfirmed metabolites were matched to numerical BinBase IDs with identical spectra and retention times. The dataset used to perform all metabolomic analyses can be found in Table S1 (https://figshare.com/s/06a7038abf59b1e5b9e2). Four animals from each group were used for these analyses.

### DNA isolation and reduced representation bisulfite sequencing

~10 mg of the left gastrocnemius was cut to contain equal amounts of both medial and lateral heads and DNA was isolated after an overnight incubation in proteinase K using the Qiagen Blood and Tissue kit following the manufacturer’s guidelines. DNA was sent to Novogene, LLC for bisulfite conversion, library preparation and reduced representation bisulfite sequencing (RRBS). DNA quality was checked with 1% agarose gel electrophoresis and quantified using a Qubit fluorometer (ThermoFisher). Samples were digested with MspI and fragments were repaired, adenylated and ligated to 5-methylcytosine-modified adapters. DNA fragments were size-selected (40 – 220 bp) using gel cutting and bisulfite treated using the Zymo Research EZ DNA Methylation Gold Kit. PCR amplification was used to generate the final DNA library. Library quality control was carried out by examining the inset size using the Agilent 2100 Bioanalyzer and the library was quantified using qPCR. Sequencing was then completed using a NovaSeq6000. DNA sequencing was completed on 4 sham animals and 6 animals for both SCIv and SCIb groups at each timepoint.

### RRBS data processing and data analysis

Paired-end fastq files generated from RRBS were checked for quality using FastQC (v0.11.9) (41), trimmed using Trim Galore (v0.6.6) and CutAdapt(v3.1) (42), and aligned against the GRCm38 reference genome using Bismark (43), which then generated methylation coverage files. From these coverage files, differentially methylated CpG sites (DMCs) were detected via methylKit [v1.16.1; (44)] in R [v4.03; (45)] and R Studio [v1.3.1093; (46)] across the different groups. HOMER motif enrichment analysis was performed on the promoter-associated (defined as CpG sites occurring in 0-5000 bp upstream of a transcriptional start site) DMCs, +/-10 bp, for each analysis using the “findMotifsGenome.pl” command (47). The R code and list of R packages used can be found in Supplementary File 1 (https://figshare.com/s/0cdef103fd8f43843bb4).

### RNA isolation and sequencing

RNA was isolated from ~20 mg of the gastrocnemius cut to contain equal amounts of both medial and lateral heads of the gastrocnemius using the miRNeasy kit (Qiagen) according to manufacturer’s recommendations, with minor deviations. Samples were homogenized in 1 ml Qiazol (Qiagen) with 5 μl β-mercaptoethanol using a beadmill homogenizer cooled with liquid nitrogen and then phase separated using 1-bromo-3-chloropropane (BCP). The clear phase was mixed with 100% ethanol and subjected to column isolation and centrifugation using buffer RWT and RPE as recommended by the manufacturer. The columns were then washed with 2 changes of 80% ethanol at 8,000 x g for 60 s at room temperature. RNA was eluted with RNase-free water heated to 37°C, quantified at 260 nm with a Nanodrop (ThermoFisher), and quality was checked with an Agilent 2100 Bioanalyzer. Samples were then shipped to Novogene, LLC for library preparation and mRNA sequencing with 1 μg of RNA per sample being used as input material for all RNA sample preparations. Sequencing libraries were generated using NEBNext UltraTM RNA Library Prep Kit for Illumina (New England BioLabs) following manufacturer’s recommendations. Briefly, mRNA was purified from total RNA using polyT, oligo-attached magnetic beads. Fragmentation was carried out using divalent cations under elevated temperature in NEBNext First Strand Synthesis Reaction Buffer (5X). First strand cDNA was synthesized using random hexamer primer and M-MuLV reverse transcriptase (RNase H minus). Second strand cDNA synthesis was then performed using DNA polymerase I and RNase H plus. Remaining overhangs were converted into blunt ends via exonuclease/polymerase activities. After adenylation of the 3’ ends of DNA fragments, NEBNext Adaptor with hairpin loop structure were ligated to prepare for hybridization. In order to select cDNA fragments of 150~200 bp, the library fragments were purified with AMPure XP system (Beckman Coulter). 3 μl USER Enzyme (New England BioLabs) was then used with size-selected, adaptor-ligated cDNA at 37° C for 15 min, which was followed by another incubation of 5 min at 95 °C before PCR. PCR was performed with Phusion High-Fidelity DNA polymerase (New England Biolabs), Universal PCR primers and Index (X) Primer. Lastly, PCR products were purified (AMPure XP system) and library quality was assessed on the Agilent Bioanalyzer 2100 system. The clustering of the index-coded samples was performed on a cBot Cluster Generation System using PE Cluster Kit cBot-HS (Illumina) according to the manufacturer’s instructions. After cluster generation, the library preparations were sequenced using a NovaSeq6000 and paired-end reads (150 bp) were generated. RNA sequencing was completed on 4 sham animals and 6 animals for both SCIv and SCIb groups at each timepoint.

### RNAseq data processing and analysis

Paired-end fastq files generated from sequencing were checked for quality using FastQC (v0.11.9) (26), then trimmed of adapters and low-quality reads (PHRED < 20) using Trim Galore (v0.6.6) and CutAdapt (42). Alignment to the mouse genome (GRCm38) and gene counts were then generated using Spliced Transcripts Alignment to a Reference [STAR; (48)]. DESeq2 was used for differentially expressed gene (DEG) analyses. DESeq2 first calculates dispersion estimates through maximum likelihood, after which it calculates gene-wise dispersion using the empirical Bayes method (49). Statistical significance was computed via a Wald test, with *p* values adjusted using Benjamini-Hochberg false discovery rate (FDR). Pathway analyses of DEG gene sets were completing using Reactome 2016, Gene Ontology: Biological Processes 2021, Wikipathway 2019, and KEGG 2019 using *EnrichR* (50). Protein-protein interaction (PPI) networks using DEG profiles as inputs were generated using STRING v11 with clustering carried out using MCODE and plotted using Cytoscape (51). Heatmaps were generated using pheatmap (v1.0.12). DESeq2 and visualization analyses were completed using R (v4.0.3) and R studio (v1.3.1093). The R code and list of R packages used can be found in Supplementary File 1 (https://figshare.com/s/0cdef103fd8f43843bb4).

We used Pathway-Level Information ExtractoR (PLIER) for exploratory RNAseq analyses (52). Read counts were filtered for low expression and normalized using trimmed mean of M values (TMM). Normalized reads were then deconvoluted using PLIER to generate latent variables (LVs) that summarize the overall structure and increase the biological interpretability of the data set. Briefly, PLIER performs a singular value decomposition constrained by prior knowledge from established pathways in the literature and other gene ontology resources. LVs were compared across samples using a mixed-model ANOVA to determine if there were any molecular patterns associated with boldine, treatment or timepoint. Data were generated using the server-hosted version of PLIER (http://gobie.csb.pitt.edu/PLIER/) with default settings, then visualized using R version 4.1.2.

### Statistics

Two-way mixed-model ANOVAs (treatment * timepoint) were used to compare body and muscle mass among groups with *p* < 0.10 being our statistical threshold for meaningful group differences, though we acknowledge there are many limitations with *p* value-based null hypothesis testing (53). Metaboanalyst 5.0 (54) was used for two-way mixed-model and one-way exploratory ANOVA analyses of metabolomics data, with an FDR <0.10 as the statistical threshold for meaningful differences. Statistical thresholds for DEGs were Ward’s *p* test with an FDR < 0.10. For DNA methylation, a chi-squared test with overdispersion correction with an FDR < 0.10 was used for principal analysis for group comparisons, with exploratory analyses completed on DMCs with raw *p* values of < 0.01 and |Δ in methylation| > 10%. The PLIER statistical threshold was conservatively set by a treatment * timepoint interaction at p = 0.05/ the number of LVs detected by the model (0.05/18), making the final PLIER threshold *p* < 0.003. Tukey’s multiple comparison tests were used for follow-up testing of LVs meeting the interaction effect threshold.

All raw fastq files for RNAseq and RRBS have been deposited to Gene Expression Omnibus under accession number GSE210392. Data files containing unnormalized read counts for RNAseq and Bismark methylation coverage files can be found using the GEO accession number and peak spectra for metabolomics can be found in Table S1 (https://figshare.com/s/06a7038abf59b1e5b9e2).

## Results

### Body mass

There was no interaction between timepoint and absolute body mass (Fig. 1A; *p*=0.901). There was a main effect of treatment on absolute body mass (*p<*0.001) with SCI groups being lower compared to sham, as well as for timepoint (*p=*0.019), with 7 d being greater than 28 d. There was no interaction effect for body mass percent change (Fig. 1B; *p*=0.187). There was a main effect of treatment (*p<*0.001), with the Sham group being greater compared to the SCI groups; there was no main effect of timepoint for percent change in body mass (*p=*0.702).

**Fig. 1.**
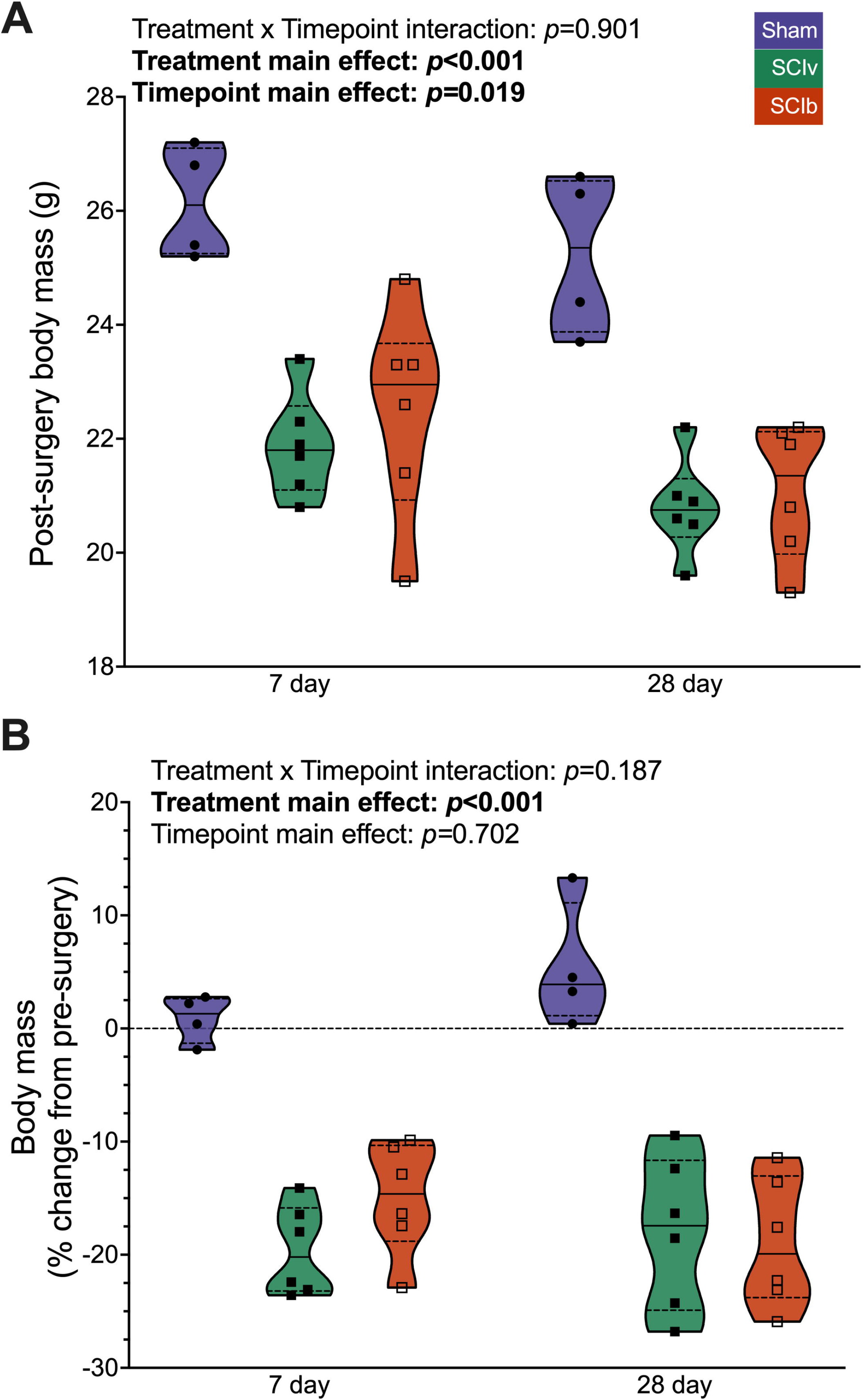
Body mass changes after SCI with and without boldine. A) Absolute and B) relative changes in body mass 7 and 28 d are reduced after a complete SCI, with no effect of boldine. Individual data points are presented in violin plots with 95% confidence interval boundaries. The solid black line represents the median value and hashed lines represent quartiles.

### Tissue mass

There was an interaction effect of timepoint and treatment for the soleus (Fig. 2A; *p<*0.001), plantaris (Fig. 2B; *p=*0.040), gastrocnemius (Fig. 2C; *p<*0.001), EDL (Fig. 2D; *p=*0.010) and TA (Fig. 2E; *p=*0.007), and. Post-test multiple comparisons showed the 28 d Sham group was elevated compared to other groups for these muscles. The 7 d soleus was the only muscle group where SCIb was elevated vs. SCIv (*p*=0.020). For the biceps, there was no interaction effect (*p*=0.631) or treatment effect (*p=*0.250). There was a main effect of timepoint (*p=*0.001), with the 28 d being greater compared to the 7 d animals (Supplemental File 2, Fig. S1A; https://figshare.com/s/3b6ae885cb2ee17b22d5). Lastly, there was no interaction effect for the triceps (*p=*0.637) or main effect for timepoint (*p=*0.454). There was a main effect of treatment (*p=*0.034), with the Sham group being elevated compared to SCI animals (Supplemental File 2, Fig. S1B; https://figshare.com/s/3b6ae885cb2ee17b22d5).

**Fig. 2.**
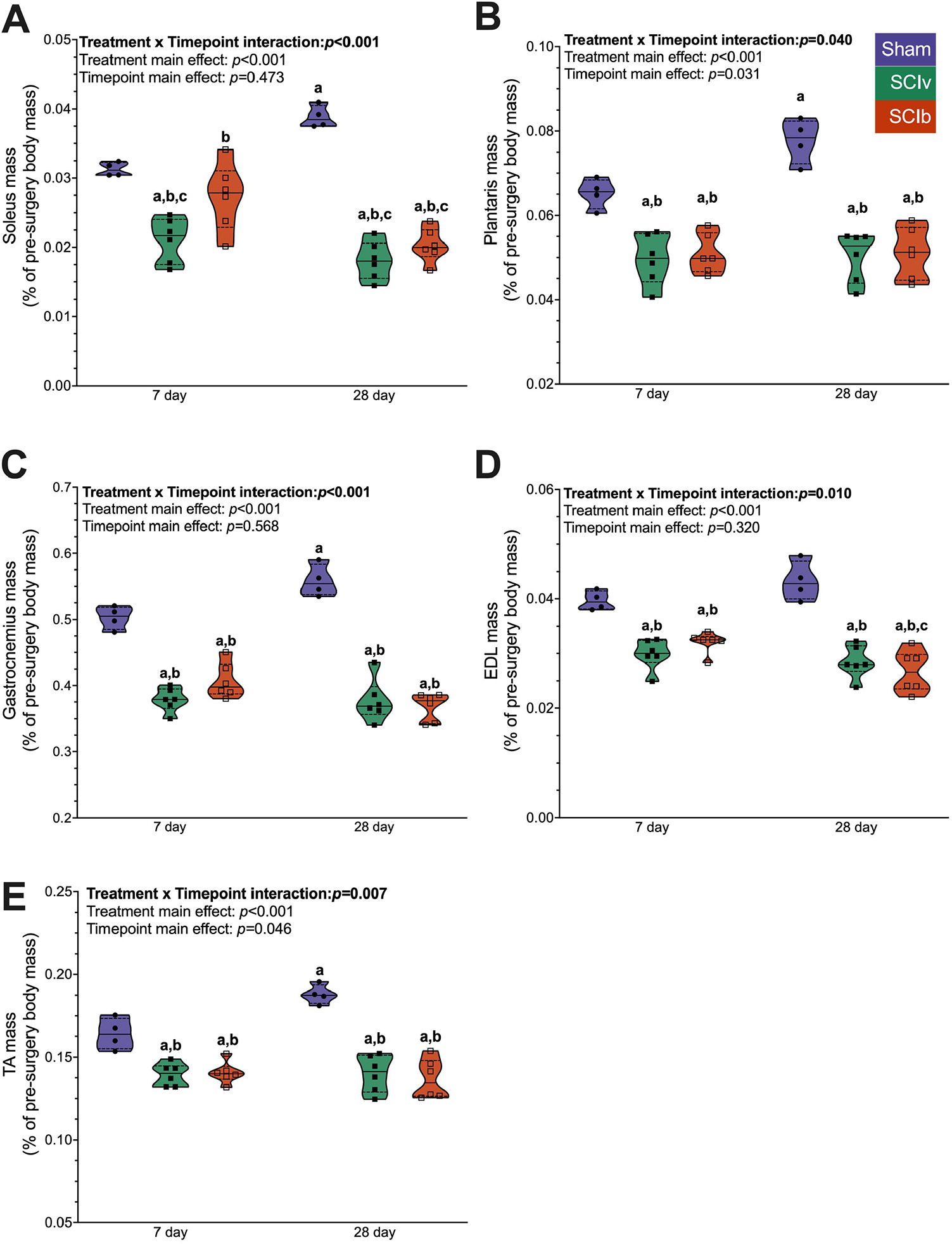
Normalized mass of the major hindlimb muscles post-injury. A) Soleus atrophy was blunted with boldine at 7 d but not 28 d post-SCI. SCI led to similar magnitudes of muscle loss with and without boldine for the B) plantaris, C) gastrocnemius, D) EDL and E) TA. Individual data points are presented in violin plots with 95% confidence interval boundaries. The solid black line represents the median value and hashed lines represent quartiles. Tukey’s multiple comparison test was used for interaction follow-up with ^a^ denoting statistical differences vs. 7 d Sham, ^b^ vs. 28 d Sham and ^c^ vs. 7 d SCIb.

### Metabolomics

575 metabolites met quality and filtering criteria so that there were no missing data points across surgeries and timepoints (n=4/group, 24 total). A PCA plot shows clustering by timepoint (Fig. 3A). Additional PCA plots show clear separation between SCIv and Sham groups, with SCIb being an intermediate cluster at 7 d (Fig. 3B, top panel). At 28 d, there was overlap among all groups (Fig. 3B, bottom panel). 22 metabolites met the criteria for an interaction effect using a mixed-model two-way ANOVA (FDR < 0.10), with 6 of them being annotated (Table S2; https://figshare.com/s/63681359100eeb32b15a). A hierarchical clustering heatmap of the 22 metabolites with an interaction effect FDR < 0.10 shows two clear interaction patterns: 1) boldine blunts the 7 d SCI response and 2) the reduced abundance of metabolites at 7 d in both SCI groups returns to sham levels by 28 d (Fig. 3C). 162 metabolites had an FDR < 0.10 for the main effect of timepoint, and 89 metabolites had an FDR < 0.10 for the main effect of treatment (Supplemental File 3; https://figshare.com/s/04384f17f5e5f603094b). Exploratory one-way ANOVAs at each timepoint resulted in 76 metabolites with FDR < 0.10 at 7 d, while only 1 metabolite had an FDR < 0.10 among groups at 28 d. At the 7 d timepoint, Tukey’s multiple comparisons follow-up testing showed boldine either prevented or attenuated SCI-induced changes in amino acids such as proline, phenylalanine, lysine, leucine and isoleucine (Fig. 3D), and other notable metabolites such as 2-hydroxyglutarate, glucose, glutathione, guanosine and inositol-4-monophosphate (Fig. 3E). The full list of differentially regulated metabolites from the one-way ANOVAs is available in Table S3 (https://figshare.com/s/a7088905168e9309adcf).

**Fig. 3.**
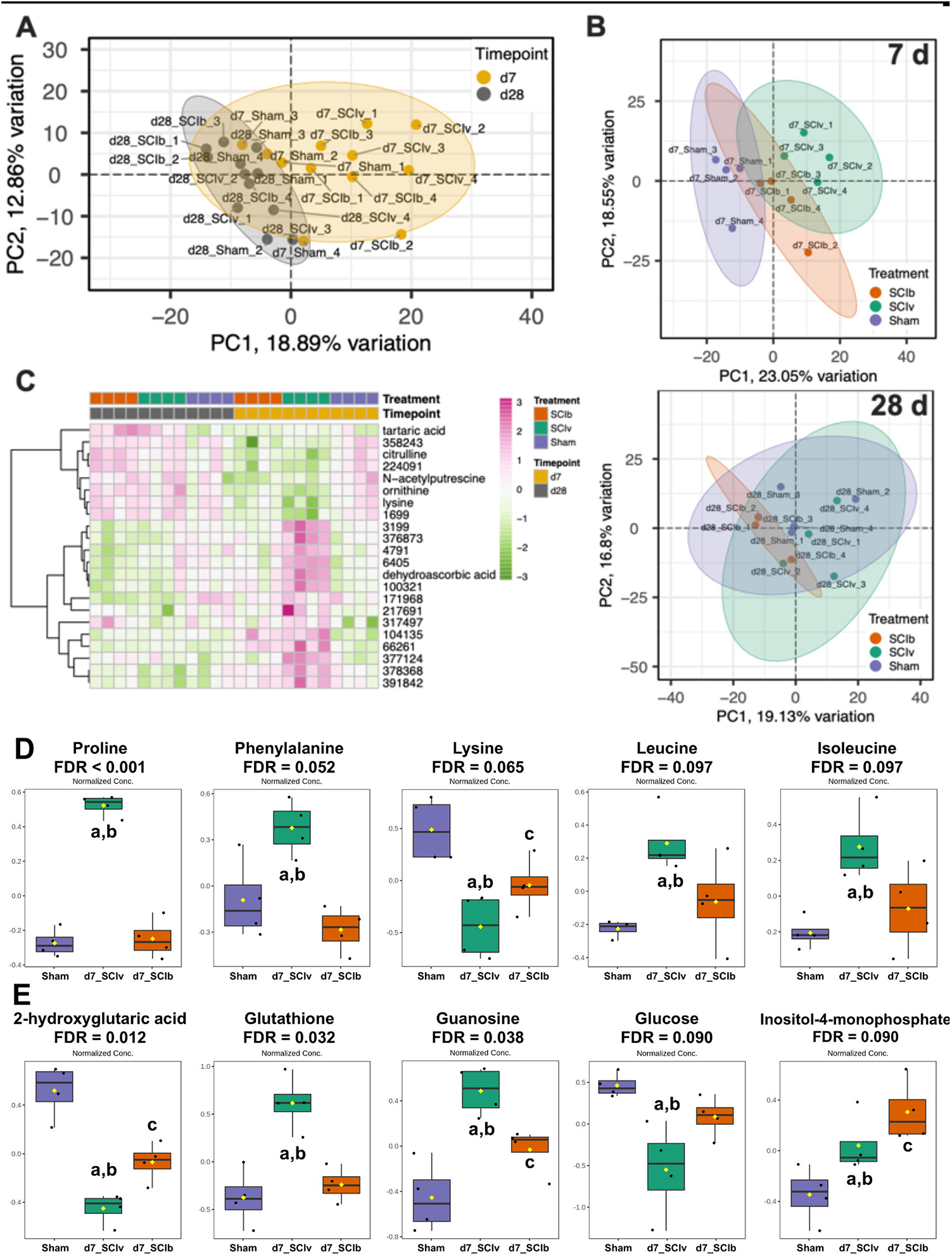
Metabolomics profiles of the gastrocnemius 7 and 28 d post-SCI. A) PCA demonstrating timepoint-based clustering of all samples. B) Top panel shows unique clustering of groups at 7 d, with the bottom panel showing more overlap. C) Fixed-sample clustering heatmap of features with FDR < 0.10 using a mixed-model ANOVA shows a clear cluster of boldine-elevated metabolites at 7 d and a cluster showing reduced expression in SCI animals at 7 d. Boldine-associated changes in annotated metabolites 7 d after complete spinal cord transection for D) key amino acids and E) other key molecules associated with muscle health. Individual data points are presented in box and whisker plots with 95% confidence interval. The solid black line represents the median value and the edges of the box represent quartiles. Tukey’s multiple comparison test was used for follow-up analyses with ^a^ denoting statistical differences vs. Sham, ^b^ vs. SCIb and ^c^ vs. SCIv

### Differential mRNA expression

We investigated the mRNA transcriptomic profiles between treatments at 7 and 28 d post-SCI separately. Our primary comparisons were ‘SCIv vs. Sham’, and ‘SCIb vs. SCIv’. Fig. 4A shows the hierarchical clustering heatmaps of DEGs across timepoints and comparisons with all three treatment groups included for relative comparison. Acute SCI resulted in the largest change in the transcriptome with 7,705 genes (3,787 upregulated, 3,918 downregulated) in the ‘SCIv vs. Sham’ comparison. ‘SCIb vs. SCIv’ led to 146 DEGs (59 upregulated, 87 downregulated) at 7 d, 105 of which were shared with ‘SCIv vs. Sham’ and 57 inversely regulated. The ‘SCIv vs.Sham’ comparison resulted in 2,046 DEGs (1,002 upregulated, 1,044 downregulated) at 28 d post-SCI, with ‘SCIb vs. SCIv’ having 110 DEGs (82 upregulated, 28 downregulated); 34 of these DEGs were shared between the comparisons. An *UpSetR* plot of the top 15 unique and shared sets of DEGs across all comparisons is presented in Fig. 4B. The ‘SCIv vs. Sham’ comparison at 7 d had ~3,000 unique up- and downregulated genes. There were DEGs shared by the ‘SCIv vs. Sham’ comparisons at both 7 and 28 d that are changed in the same direction (685 downregulated, 477 upregulated), with additional sets of up- and downregulated genes specific to SCI at the 28 d timepoint. Other intersections of note show 36 genes unique to upregulation in the ‘SCIb vs. SCIv’ comparison at 28 d, as well as 31 genes shared in the 7 d ‘SCIb vs. SCIv’ upregulated comparison that were downregulated in the 7 d ‘SCIv vs. Sham’ comparison.

**Fig. 4.**
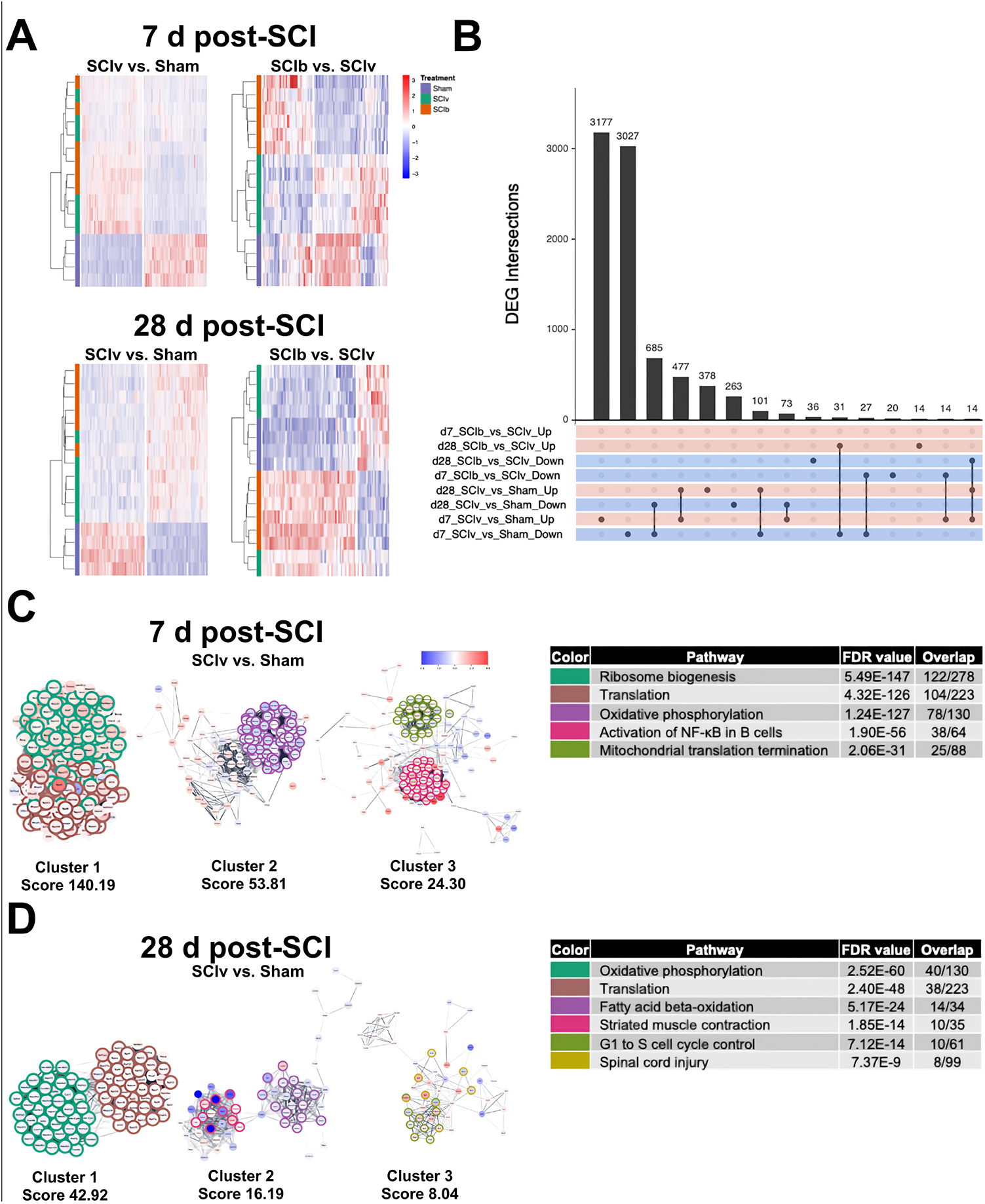
Transcriptomic signatures of complete SCI at acute and subacute timeframes after injury. A) Hierarchical clustering heatmaps across the main comparisons and timepoints. B*) UpSetR* plot showing unique and overlapping DEG sets across all comparisons. C) STRING association networks and related MCODE cluster pathway analyses showing clear clustering of key biological processes expected to be affected by SCI at both 7 and D) 28 d post-injury.

### Pathway analyses, network clustering and PLIER

*EnrichR* was used to perform pathway analyses on the DEGs from each comparison. The top 5 non-redundant pathways and ontologies for both upregulated and downregulated gene sets are shown in Table 1. PPI networks were generated for each analysis with the goal of identifying clusters of highly related genes and their associations with specific pathways. MCODE analysis within the PPI networks identified clusters from the 7 d ‘SCIv vs. Sham’ comparison showing cluster 1 as upregulated genes associated with ‘Ribosome biogenesis’ and ‘Translation’, with cluster 2 demonstrating downregulation of genes associated with ‘Oxidative phosphorylation’. Cluster 3 highlights downregulation of DEGs associated with ‘Mitochondrial translation termination’ and upregulation of genes annotated to ‘Activation of NF-κB in B cells’ (Fig. 4C). The top 3 MCODE clusters and cluster specific pathways for the ‘SCIb vs. SCIv’ comparison are shown in Supplemental File 2, Fig. S2A (https://figshare.com/s/3b6ae885cb2ee17b22d5). Each of these clusters had only 3-5 genes per comparison. Cluster 1 of the 7 d ‘SCIb vs. SCIv’ comparison was annotated to ‘Ubiquitin-mediated proteolysis’, cluster 2 to ‘Starch and sucrose metabolism’ and cluster 3 to ‘N-glycan trimming in the ER and calnexin/calreticulin pathway’. The top clusters from the ‘SCIv vs. Sham’ comparison at 28 d showed maintained alterations in pathways annotated to ‘Oxidative phosphorylation’ and ‘Translation’ for cluster 1. Cluster 2 was composed of downregulated genes enriched for ‘Fatty acid oxidation’ and ‘Striated muscle contraction’. Cluster 3 was composed of DEGs that annotated to ‘G1 to S cell cycle control’ and ‘Spinal cord injury’ (Fig. 4D). Cluster 2 was annotated to ‘Focal Adhesion’ for ‘SCIb vs. SCIv’ at 28 d; no other cluster met FDR criteria (Supplemental File 2, Fig. S2B; https://figshare.com/s/3b6ae885cb2ee17b22d5). The full list of significant pathways be found in Supplemental File 4 (https://figshare.com/s/d781f7cf7e185410c595).

**Table 1.**
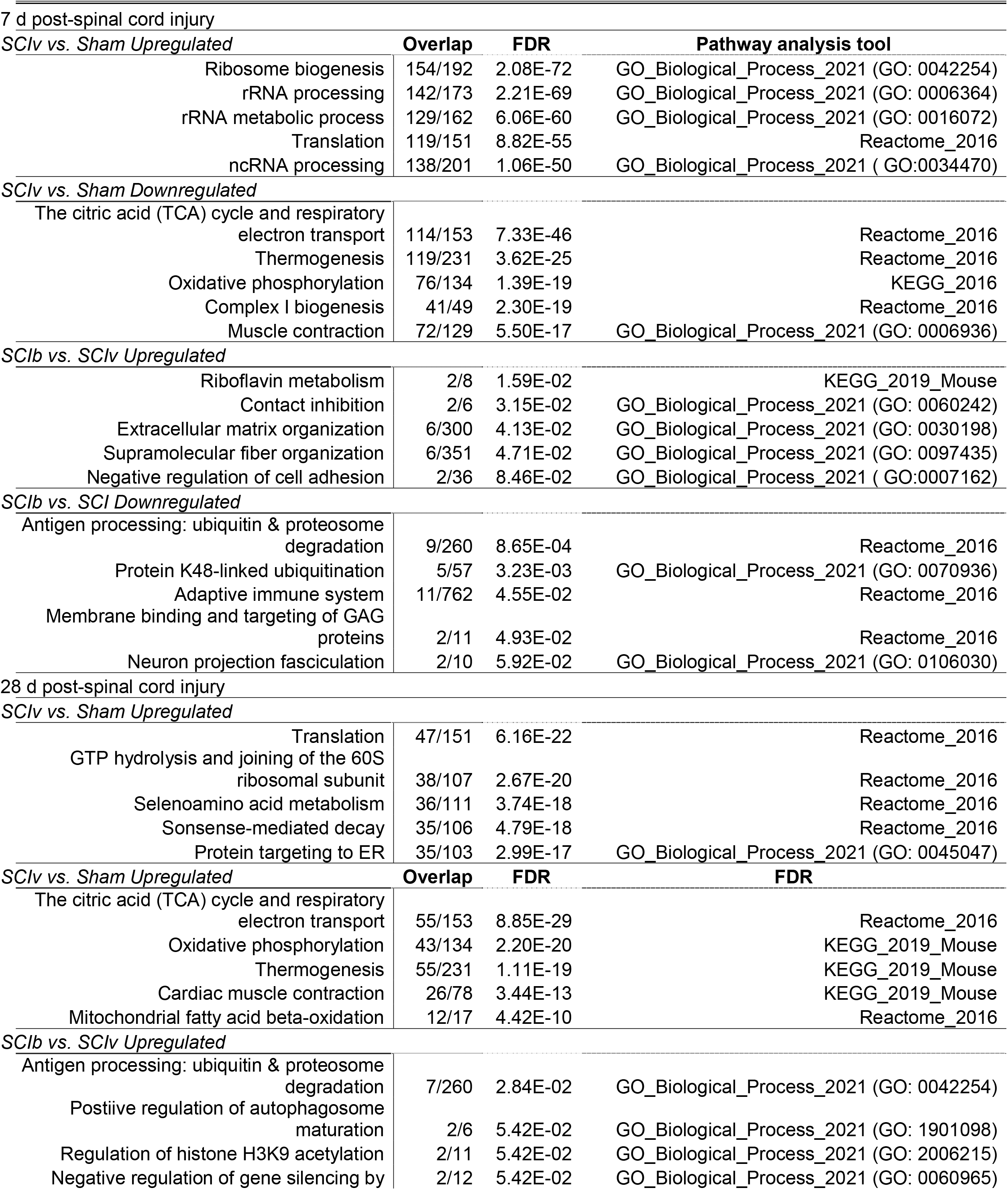
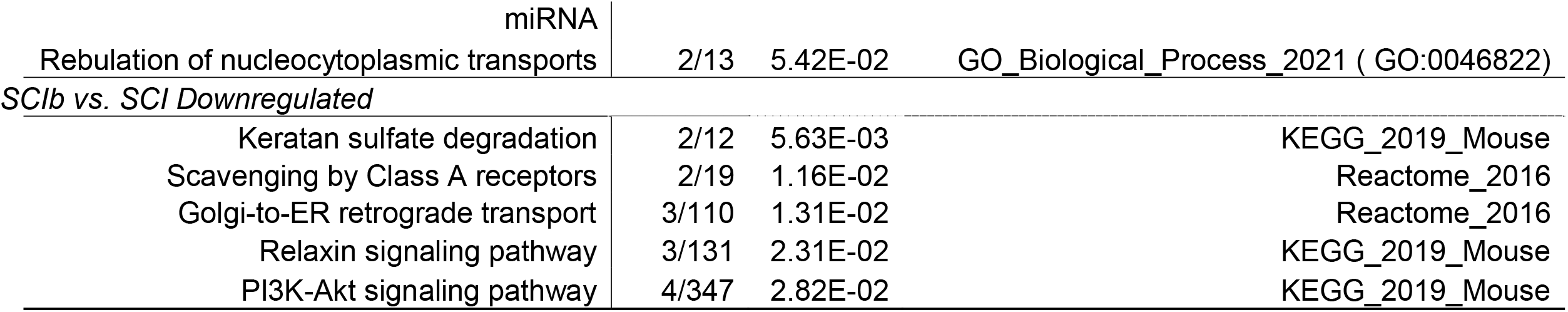
Selected top pathways enriched after spinal cord injury.

PLIER constructed 18 LVs following deconvolution (Table 2). Three LVs had an interaction effect for timepoint and treatment (LV1, 3 and 5), 5 had a main effect of timepoint (LV1, 2, 3, 5 and 7) and 4 had a main effect of treatment (LV1, 2, 3 and 5). The LVs constructed by PLIER contained genes associated with pathways identified above such as alterations in translation and mitochondrial ribosomes (LV1, 2, 5). Heatmaps of the top-loaded genes for these constructed LVs and box plots of the individual sample scores highlighting the interaction effects are shown in Figs. 5A and 5B, respectively. LV1, annotated to ‘MIPS Nop56p-associated pre-rRNA Complex’ and LV5, annotated to ‘MIPS Spliceosome’, demonstrated clear SCI-induced timepoint effects without any effect of boldine. LV3 annotated to ‘PID Androgen Receptor Transcription Factor Pathway’ and showed a potent effect of the Sham surgery at 7 and 28 d. In addition, it showed a unique boldine effect as group loading values were elevated in the SCIb animals compared to SCIv at 28 d. LV2, annotated to ‘MIPS 55S Ribosome Mitochondrial’ and LV7, annotated to ‘Reactome Antigen Processing Ubiquitination and Proteasomal Degradation’, are presented in Supplemental File 2, Fig. S3. LV2 had strong SCI-induced downregulation at 7 d, with all groups having a positive response by 28 d. LV7 had SCI-induced upregulation at 7 d, with all treatment groups having similar values by 28 d. All multiple comparison follow-up test *p* values can be found in Table S5 (https://figshare.com/s/4c1edaf591d9e999afc2).

**Table 2.**
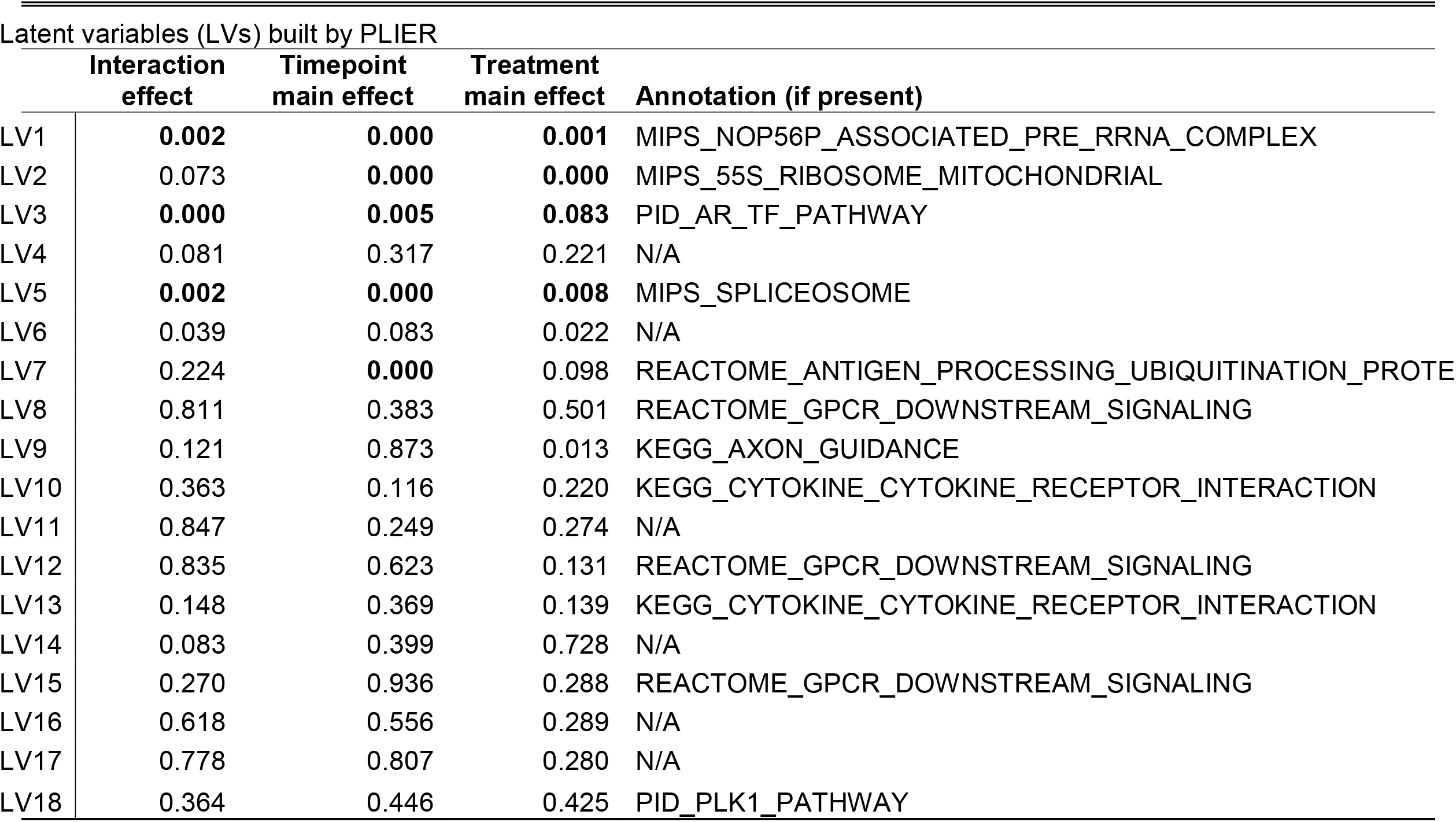
Pathway Level Information ExtractoR (PLIER) Results.

**Fig. 5.**
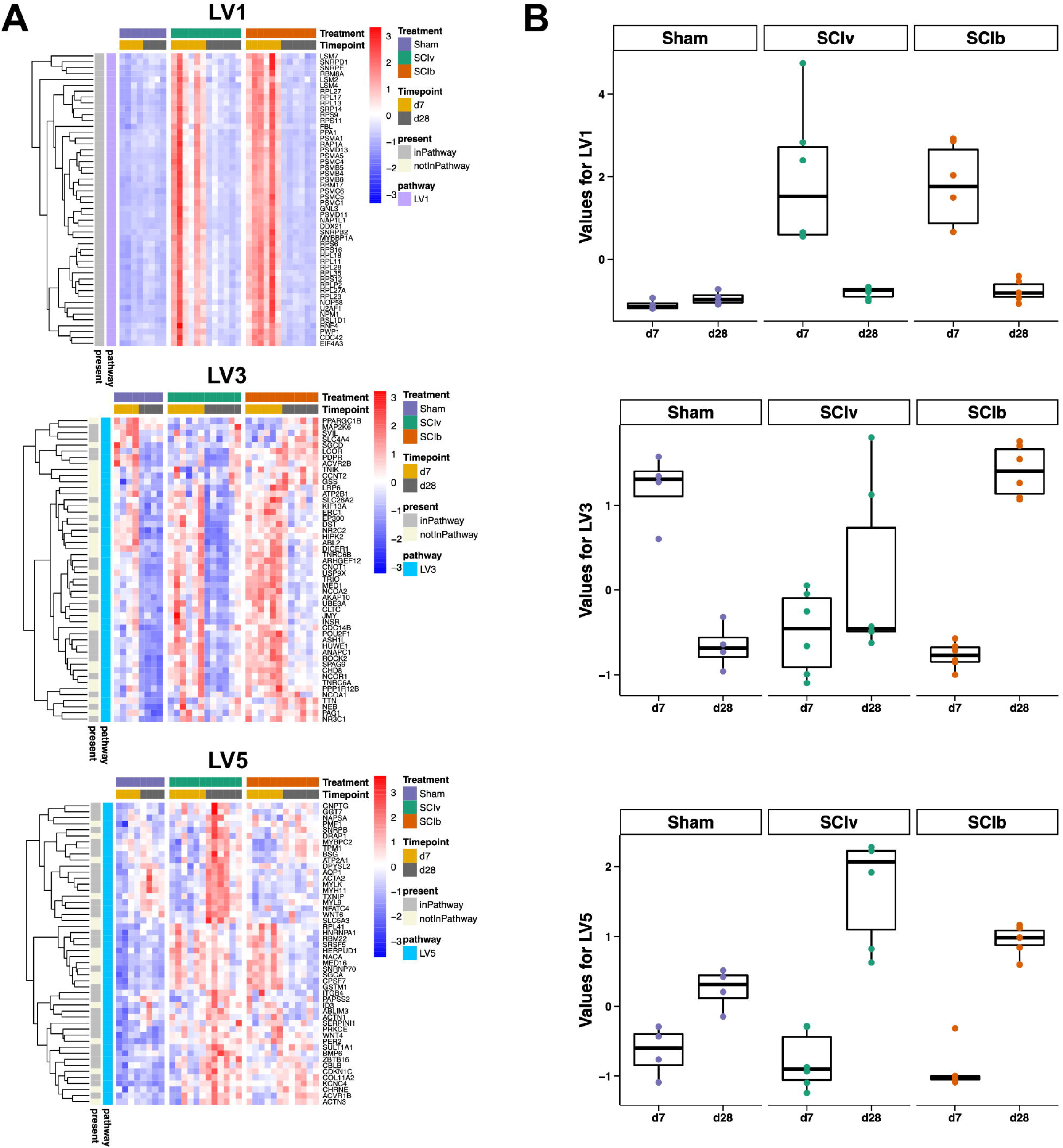
Pathway Level Information ExtractoR (PLIER) highlights unique molecular circuits after SCI. A) Heatmap of the top loaded genes from constructed LVs with timepoint * treatment interaction effects. B) Box plots showing the overall loading score of each animal for the respective LV.

### DNA methylation

The RRBS analysis strategy was identical to that used for RNAseq, with analyses separated by timepoint and key comparisons being ‘SCIv vs. Sham’ and ‘SCIb vs. SCIv’. An FDR < 0.10 resulted in < 10 DMCs in each comparison at both timepoints (see Table S6). We thus expanded our approach using DMCs with a raw *p* value < 0.01 and absolute methylation change > 10%. The ‘7 d SCIv vs. Sham’ comparison resulted in 2,991 DMCs (1,614 hypermethylated, 1,377 hypomethylated) across 1,781 annotated genes, with the ‘SCIb vs. SCIv’ comparison resulting in 2,217 DMCs (1,199 hypermethylated, 1,018 hypomethylated) across 1,365 genes. Similar numbers of DMCs were observed at 28 d, with heatmaps showing DMC profiles for each comparison in Fig. 6A. SCI resulted in 2,645 DMCs (872 hypermethylated, 1,773 hypomethylated) across 1,560 genes compared to sham animals, with boldine treatment leading to 2,459 DMCs (1,742 hypermethylated, 717 hypomethylated) across 1,419 genes compared to vehicle after SCI. DMC intersections among comparisons are shown in Fig. 6B. The *UpSetR* plot shows each comparison had a unique set DMCs. There were also 129 DMCs that were inversely methylated at 28 d and 160 inversely methylated DMCs at 7 d. The complete outcomes for all DMCs are provided in Supplemental File 5 (https://figshare.com/s/af57bff7bdf3f5dc6a45).

**Fig. 6.**
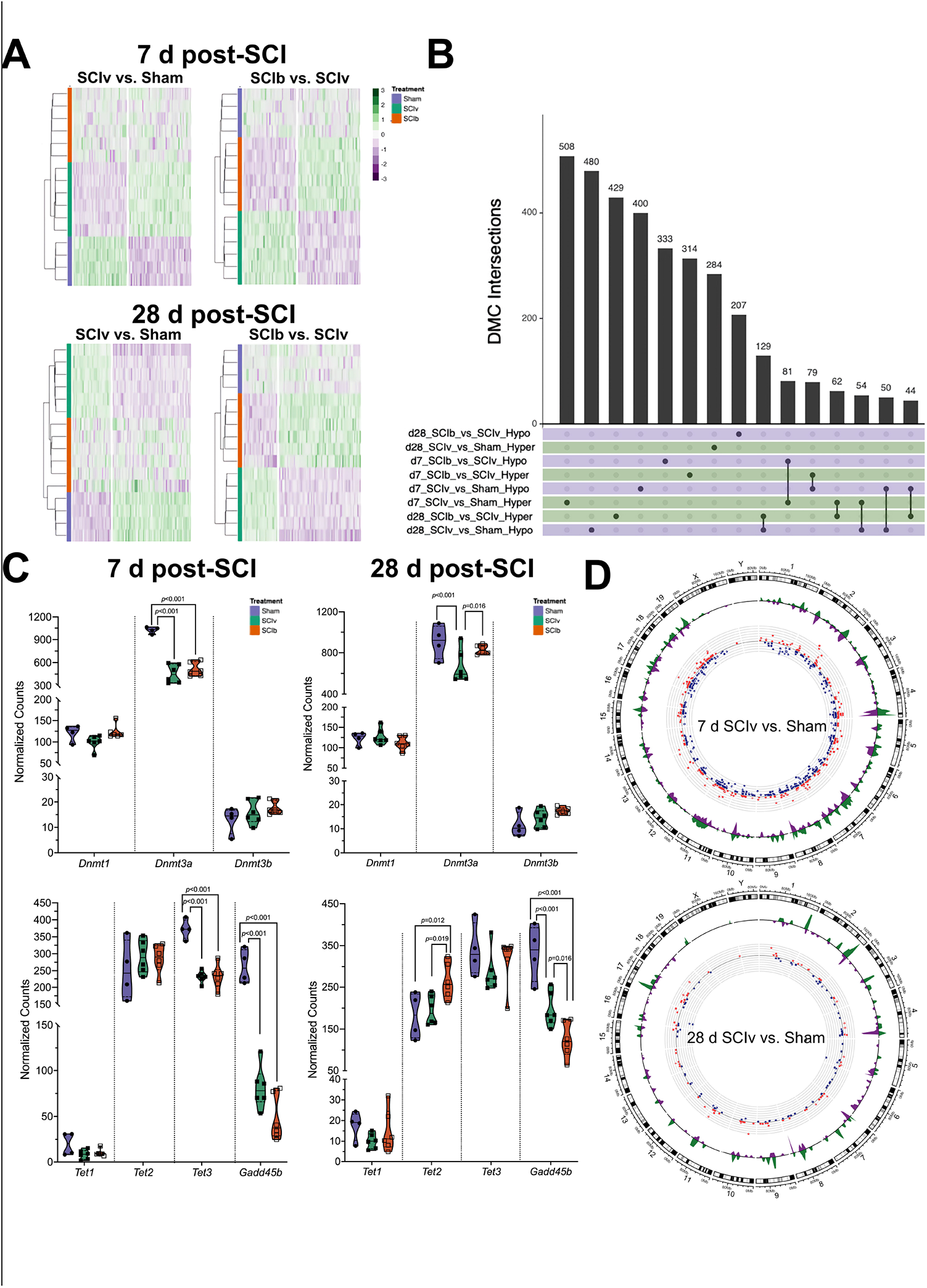
DNA methylation of the gastrocnemius post-complete SCI. A) Hierarchical clustering heatmaps of the most hyper- and hypomethylated genes for each comparison. B) *UpSetR* plot showing that each comparison has a mostly unique set of DMCs, with the shared DMCs for ‘SCIb vs. SCIv’ and ‘SCIv vs. Sham’ comparisons at both 7 and 28 d being inverse to each. C) gene expression using normalized read counts of factors associated with methylation and demethylation. *Dmt3a, Tet3 and Gadd45b* are greatly downregulated at 7 d, with boldine being associated with improved levels of *Dmt3a* and reduced expression of *Gadd45b* at 28 d. D) Circos plots demonstrating the overlap and magnitude of change in DEGs and DMCs across the ‘SCIv and Sham’ comparisons at 7 and 28 d post-SCI. Red dots = upregulated genes, blue dots = downregulated genes; green waves = hypermethylated DMCs, purple waves = hypomethylated DMCs.

Promoter-associated CpG sites, split by hyper or hypomethylation, were run through HOMER motif enrichment analysis to identify differences in methylation of key transcription factor binding motifs. The results are presented in Table S5 (https://figshare.com/s/71fd8345296834deb3da). The 7 d ‘SCIv vs. Sham’ comparison identified hypermethylation of the motif for PRDM1. The ‘SCIb vs. SCIv’ comparison resulted in no hypermethylation that met FDR criteria, but was associated with hypomethylation of motifs for ELK1 and 4, ETV 1 and 4, FLI1, SPDEF and EFL1 (FDR < 0.10). The ‘SCIv vs. Sham’ comparison at 28 d resulted in no motif meeting hyper-or hypomethylation FDR criteria. There was no hypermethylation of motifs associated with the ‘SCIb vs. SCIv’ comparison at 28 d but hypomethylation of ELF1.

The 7 d timepoint resulted in differential regional proportions of DMCs (Supplemental File 1, Fig. S4). Regional DMCs were roughly the same 1 to 5 kb upstream (~10%), in the promoter region (~5%) and in the 5’ UTR (~1-2%). However, the ‘SCIv vs. Sham’ comparison resulted in more DMCs within exons (~17% vs. 2%), intron/exon boundaries (~15% vs. 7%) and introns (40% vs. 15%). The ‘SCIb vs. SCIv’ resulted in more DMC proportions in the 3’ UTR (~38% vs. 2%) and intergenic regions (~16% vs. 7%). The regional DMC proportions were identical between the comparisons at 28 d post-SCI.

Normalized read counts from our transcriptomics data shows the major DNA methyltransferase in skeletal muscle, *Dnmt3a*, reduced by SCI with no effect of boldine (Fig. 6C, top left panel). Normalized read counts of genes associated with DNA demethylation shows *Tet3* and *Gadd45b* reduced following SCI, with no change in *Tet2* and minimal overall expression of *Tet1* (Fig. 6C, bottom left lower panel). Expression of *Dnmt3a* was preserved with boldine treatment at 28d using normalized read counts, with boldine leading to increased expression of *Tet2* and further reductions in *Gadd45b* (Fig. 6C, top and bottom right panels, respectively).

### DEG and DMC overlap

We plotted the overlap of DEGs and DMCs for each comparison. The 7 d ‘SCIv vs. Sham’ had 658 DEGs that overlapped with its altered methylation status (Fig. 6E, left panel), with this number reduced to 165 at 28 d (Fig. 6D). The ‘SCIb vs. SCIv’ comparison resulted in 11 overlapping genes at 7 d ‘SCIb vs. SCIv’ and 8 at 28 d (Supplemental File 2, Fig. S2).

## Discussion

SCI results in rapid muscle loss and drug/small molecule interventions have largely been ineffective in slowing these losses following severe contusion and complete transection SCI. In the current report, we show boldine treatment was not able to slow or prevent body and hindlimb muscle mass losses at 7 or 28 d after a complete thoracic spinal cord transection. However, boldine prevented or blunted changes in abundance of glucose and amino acids 7 d post-SCI. Boldine also altered expression of a select group of mRNAs and resulted in unique DMC signatures. These molecular profiles provide an attractive starting point for future studies to determine the biological significance of CxHC on skeletal muscle health and function using other SCI or atrophy models.

This study did not show preservation of body or tissue mass after SCI with boldine, with the exception of the soleus at 7 d post-SCI, suggesting CxHCs are not a primary driver of muscle atrophy after SCI. Muscle requires loading to maintain mass and function, and the amount of loading required is surprisingly minimal. For example, 60 maximal isometric contractions per day can preserve muscle mass and specific tension in paralyzed rats 28 d after a spinal cord isolation, a surgical procedure that creates an almost electrically silent muscle via spinal cord transection at two levels as well as bilateral dorsal rhizotomy between transection sites (14, 55). Moderate to moderate-severe SCI (~50-80 kdyne injury force in mice) does not result in complete paralysis (56–59). This potential for reloading of muscles in combination with pharmaceuticals has been recently shown to have some efficacy in improving muscle mass and locomotor recovery (57–60). Thus, any appreciable muscle preservation using boldine would likely have to be in combination with some form of loading. The reduced severity of muscle loss in the soleus is intriguing as it is the most oxidative of the hindlimb muscles in mice, and CxHC seem to reappear de novo only in denervated fast twitch fibers (27). That same report showed evidence that complete SCI results in elevated sarcolemmal expression of Cx39, 43 and 45 in the gastrocnemius 56 d after the injury. A more recent report has shown sarcolemmal CxHC appearance does not occur with active acetylcholine receptor signaling at the neuromuscular junction (30), suggesting that anatomical or functional denervation is a major cue for CxHC formation in skeletal muscle. This interpretation is somewhat complicated as our gastrocnemius transcriptomics data shows downregulation of *Gja1* and *Gjc1*, the genes that encode Cx43 and 45, respectively, at 7 d post-SCI. Further, there is very little known regarding the role of SCI on denervation and motor unit loss in the acute and subacute timespans post-injury. There is an approximate ~30% loss in quadriceps motor neuron counts 6 months following an L2 contusion SCI in rats (61) and motor neuron death is highest at the level just below the epicenter of injury (62). The transection used in our currently study was completed at a low thoracic level and it is quite possible there is very limited motor neuron loss in our timespan as the mouse gastrocnemius is innervated by motor neurons from T13-L1 (63). Our RNAseq data is suggestive of at least neuromuscular junction remodeling as *Musk, Ncam1* and *Scn5a*, common markers of denervation, are upregulated in the SCIv animals compared to sham at 7 d, with *Musk* and *Ncam1* remaining elevated at 28 d. However, motor unit loss is not entirely indicative of denervation as nearby motor neurons can sprout axons to ‘rescue’ denervated fibers (64). A potentially simpler explanation may be a pre-clinical rat model may not be fully recapitulated in a pre-clinical mouse model. In addition, the boldine intervention began 3 d after surgery when the molecular program of muscle atrophy after SCI is at or near its peak but muscle mass has not greatly changed (65). That this preservation is not maintained at 28 d is not surprising due the severity of a complete spinal cord transection, but the observation of at least partially slowed atrophy suggests a potential for preferential oxidative fiber-type preservation by boldine.

Molecular profiling of the gastrocnemius showed intriguing outcomes despite the degree of muscle loss. For metabolomics, the 7 d timepoint was the most affected by complete SCI, similar to our previous report in female mice (10). Boldine was able to shift the overall PCA profile towards that of sham animals at 7 d. Our mixed-model approach demonstrated a unique cluster of metabolites protected by boldine at 7 d. Dehydroascorbic acid was the only annotated metabolite in this cluster. Its reduction in the presence of boldine, which also occurred with glutathione abundance at 7 d in our exploratory analyses, supports boldine’s role in preventing accumulation of oxidative species, likely via inhibiting Ca ^2+^ influx and the initiation of cell signaling pathways involved in generating oxidative stress. We demonstrated SCI induced changes in amino acids, which may be expected post-SCI due to protein degradation and anaplerotic potential (66). Why these amino acids were affected by boldine is unclear. Our transcriptomics data shows boldine was able to downregulate genes associated with protein ubiquitination at 7 d including *Ubb*, which encodes ubiquitin itself, genes associated with the ubiquitination process such as *Vps37a, Cul3, Asb11/14* and *Ube2e1*, as well as multiple E3 ligases (*Arih2, Amfr, Nedd4l, Herc3* and *Wwp1*). Thus, one possibility is reduced protein breakdown through reduced ubiquitination, though this was not sufficient to prevent atrophy as measured by muscle weights. Another possibility is protection against ECM turnover. Proline is a major component of collagen, the most abundant ECM protein in the skeletal muscle environment (67). Matrix metalloproteinases are the predominant enzymes for protein breakdown in the ECM and are activated by cellular processes that can be regulated by boldine such as inflammation and oxidative stress (68); boldine reversed expression of *Adam9*, a widely expressed metalloprotease upregulated in pathological conditions (69). Boldine also upregulated *Pi16*, which encodes peptidase inhibitor 16 (PI16), a predicted negative regulator of peptidase activity in the ECM. PI16 has been localized to the cell membrane, upregulated in diseased cardiac tissue and may be an indicator of cardiac stress (70). Similarly, genes associated with the extracellular matrix, *Postn, Col12a1* and *Col14a1* were upregulated in the SCIb animals compared to SCIv, as well as *Pxdn*, which encodes peroxidasin, a protein involved in extracellular matrix formation. Thus, an additional potential mechanism of action for boldine may be inhibiting the breakdown of proline-rich ECM proteins like collagen through upregulation of PI16. Unfortunately, 4-hydroxyproline was not detected, or failed to be confidently annotated, in our metabolomics dataset. This limits stronger support for collagen turnover and ECM remodeling as 4-hydroxyproline is a specific collagen post-translational modification.

Skeletal muscle phenylalanine release is increased during catabolism from fasting and severe burn (71, 72) but how boldine may alter this is unknown. The effect of boldine on lysine abundance is also an interesting outcome. Lysine can prevent the degradation of myofibrillar proteins through mTOR, Akt and autophagic mechanisms (73, 74). We have additional evidence of the potential for preserved anti-catabolic signaling with boldine. The SCIv animals had reduced glucose levels similar to our previous report that also noted reduced lactate and pyruvate (10), though this current study did not identify these glucose metabolites. Boldine prevented the SCI-induced reduction in muscle glucose abundance. SCI resulted in elevated *Ide* expression, which encodes insulin degrading enyzme (IDE), and boldine was able to prevent this elevation, suggesting the potential for longer duration of insulin action. Lastly, the levels of inositol-4-monophosphate were elevated in the SCIb animals compared to both SCIv and sham animals. Cellular levels of inositol and related metabolites are usually inversely regulated by glucose levels (as seen in the SCIv and Sham animals), but they can act as insulin mimetics once in the cell via second messenger systems (75). While admittedly speculative, our cumulative data hints at boldine being able to transiently preserve glucose and amino acid levels through insulin signaling pathways that both increase glucose transport and activate anti-catabolic processes during acute SCI. A systematic study of boldine’s role in regulating intramuscular glucose would be an important future direction.

Acute SCI led to major disruptions in gene expression between SCIv animals and sham controls. While SCI likely has distinct mechanisms that result in a unique form of chronic muscle atrophy due to the interaction of the spinal lesion and innervating motor neurons, the molecules involved in the immediate days post-injury share pathways common to other models of disuse atrophy. Peripheral nerve crush results in ~7,000 DEGs in muscle 3 d after injury, with gene set enrichment and pathways analyses associated with the TCA cycle, protein translation and protein synthesis (76). mRNA profiles from mice that were fasted, casted or denervated led to upregulation in genes associated with ribosomal processing and protein catabolism observed alongside the downregulation of genes associated with metabolism and the extracellular matrix (77). These principal pathways are shared in our study, highlighting that SCI-induced atrophy follows the general atrophy program at the transcriptomic level.

Transcriptomics analysis packages like *DESeq2, edgeR*, and *limma* can be used for complex dataset analysis by designating a reference group and using general linear models to determine interaction and main effects. A limitation to this is the conservativeness of statistical outcomes from the number of multiple comparisons required across the number of groups and levels in the study design. PLIER offers a simpler but distinct approach by using singular value decomposition for unsupervised deconvolution of the dataset, which is then mapped to prior information contained in available pathway and ontological databases (52). This unsupervised deconvolution can also construct LVs that do not map to known databases to provide novel molecular relationships. Additionally, the LVs generated by PLIER reduce the dimensionality of the data and permit the use of traditional multivariate testing, similar to other outcomes in this extensive study. PLIER found SCI-induced changes in translation and mitochondrial function (LVs 1, 2, 5 and 7), similar to the pathways analyses noted above. LV2 and LV7 highlight similar temporal responses to surgery, as LV2 was lower in groups at 7 d vs. 28 d and LV7 was higher at 7 d compared to 28 d. Both of these LV responses were annotated to relevant pathways: mitochondrial ribosome function (LV2) would be expected to be compromised after a severe surgery such as a laminectomy and further affected by acute paralysis. Inversely, LV7, which annotated to ubiquitination and proteasomal processes, would be expected to be elevated at 7 d post-SCI then abate in strength over time. However, the main objective with the use of PLIER was to find LVs altered in SCIb animals compared to SCIv animals. PLIER found one unique molecular signature for boldine, LV3, which was upregulated in the 28 d SCIb group compared to 28 d SCIv and annotated to the GSEA ‘PID Androgen Receptor Transcription Factor Pathway’. This gene set is involved in the regulation of androgen receptor activity. Boldine has not been linked to these cell processes in any tissue to our knowledge. Hypogonadism is a known secondary medical outcome after SCI. Severe SCI reduces serum levels of testosterone 28 d post-injury in male rats (78) and cross-sectional studies have shown the proportion of those with low total and free testosterone is 4-5 fold higher in individuals with SCI compared to able-bodied controls across the lifespan (79, 80). Thus, while the data are intriguing, more rigorous follow-up is needed to determine how androgen signaling may be altered and whether it is a function of improved serum testosterone levels, intracellular signaling or other potential mechanism. Additionally, LV3 provides interesting information regarding the laminectomy control surgeries, as there were clear time-related group differences between the 7 and 28 d Sham animals. This demonstrates the importance of having sham surgical controls even for analyses of tissues peripheral to a surgical site.

Data focused on the methylome of acute and subacute SCI do not exist to our knowledge, though array data do exist for individuals with chronic SCI (81, 82). Surprisingly, few of our data met traditional FDR thresholds for differential methylation across comparisons and timepoints. RRBS has been used to show changes to the methylome of mouse gastrocnemius in aging and exercise studies using more conservative FDR criteria than our report (83) as well as in related mouse models using exercise to investigate methylation changes in myonuclei and interstitial nuclei (84). The inter-mouse heterogeneity in response to either a laminectomy, which in itself is a severe surgery, or a spinal cord transection could lead to sufficient variability in detected methylation as to reduce the ability of our experiment to detect significant methylation changes. For example, methylation of a DMC in *Acsl5* at 28 d had a 29% mean difference between SCIv and Sham groups. The range of methylation in the SCIv animals was 45-56% in 5 out of 6 animals and 29% in the other one. The sham animals ranged from 17-23%. The variability caused by that one SCIv mouse resulted in an FDR = 0.27. Additionally, while a complete spinal cord transection is a uniform model of paralysis, it is possible that the rates of spinal cord and motor neuron degeneration were sufficiently different among animals to drive variation in detected methylation.

We used less conservative statistical thresholds for the methylome for exploratory analyses, as the reduced read counts we observed for *Dmnt3a, Tet3* and *Gadd45b* across time suggested the potential for differences in methylation profiles. Our DMC *UpSetR* plot showed the largest intersections were distinct to each comparison. However, boldine was able to revert the methylation state of 129 DMCs at 28 d and 160 at 7 d post-SCI. We then investigated whether there were changes in transcription factor motif methylation state using HOMER. PRDM15 was detected in the 7 d ‘SCIv vs. Sham’ comparison and though it has not been described in skeletal muscle, it has been shown to regulate the PI3K signaling cascade in cancer cells to improve metabolic function (85). Multiple hypomethylated motifs related to erythroblast transformation specific (ETS) family members met FDR criteria in the 7 d ‘SCIb vs. SCIv’ comparison. Due to the range of overlap of these factors, it is difficult to highlight a key biological process that would be related to boldine administration. Similarly, no strong candidate motif was detected for any comparison at the 28 d timepoint. We do provide a consistent pattern of mRNA changes that are associated with the methylation state of their gene in the ‘SCIv vs. Sham’ comparisons at both 7 and 28 d. This is not seen in the ‘SCIb vs. SCIv’ comparisons due to the relatively small number of DEGs compared to DMCs. These data provide a nice summation of our study but since we did not probe chromatin accessibility or quantify DMRs within key promoter and other regulatory sites, we cannot infer key biological meaning from the overlap alone.

Our study has several limitations: 1) This study only investigated male mice. In humans, males make up ~80% of SCI (86), which is even further skewed (~95%) within the Veteran population (87). These data may not adequately reflect what occurs in female mice though portions of our metabolomics data support our previous investigation in female mice. A follow-up study repeating these approaches using female mice would be a great complement to our report; 2) We used a complete spinal cord transection because it results in a repeatable and consistent model of paralysis. However, this is not a direct clinical analog as anatomically complete transections are rare in humans. Studies investigating these parameters in various models of contusion SCI would be a great benefit to the field; 3) The unexpectedly low number of FDRs for methylomics greatly hinders rigorous biological interrogation. While our two parameter exploratory analyses using raw *p* values and a +/-10% methylation change likely captures legitimate physiological changes, we understand that this increases the chance for a large number of potential false negatives. Further investigations using high-throughput arrays or other methylation library preparations (e.g., TWIST), as well as alternative bioinformatic pipelines, may help increase the rigor for these types of analyses; 4) Boldine may target cellular processes not directly related to CxHCs. For example, pannexins can act similarly to CxHC by allowing small molecules such as calcium into the cell (26, 31), although muscle-restricted knockout does not prevent atrophy after denervation like CxHCs (27); 5) We solely focused on skeletal muscle in this report. Our intervention was systemic so there is a chance any changes we have observed may be coming from temporally preserved function of upstream factors such as the motor neuron or even the spinal cord itself; and 6) Changes in single DMCs do not necessarily account for changes in mRNA expression. Chromatin profiling and accessibility using ATACseq would be a very useful complementary approach, as accessibility changes surrounding a DMC (or group of nearby DMCs) would imply biological significance.

In summary, SCI resulted in a severe alteration in the multiome with the most drastic changes observed during the acute timeframe. Boldine was ineffective in preserving body and muscle mass after injury though it was able to alter the metabolome at 7 d with provocative findings related to amino acids, energy substrates and multiple unannotated metabolites. Boldine had a unique transcriptomic signature related to molecular regulation of glucose metabolism and the ECM. Analyses of the methylome highlighted unique DMCs across comparisons. Our data provides a foundation for better understanding the changes that occur in paralyzed muscle in the acute and subacute timeframe after a complete SCI and provides a deeper understanding of the effects of boldine on skeletal muscle in SCI.

## Acknowledgements

This study was funded by a Department of Veterans Affairs Office of Research and Development RR&D Service CDA-2 grant (1IK2RX002781 to ZAG) and Center grant (5I50RX002020; PI, William Bauman). LAP is supported by a National Institutes of Health F31 Award (F31HL154571). Metabolomics services were completed by West Coast Metabolomics (UC2ES030158; PI, Oliver Fiehn). The views represented in this manuscript are not reflective of the United States Government or the Department of Veterans Affairs.

## Disclosures

Authors ZAG, CAT and CPC are co-inventors of a submitted patent application for the use of boldine to treat neurological injuries. This study falls within the claims of the patent.

## Author Contributions

CAT, CPC and ZAG conceived and designed research. CAT, LH and ZAG performed experiments. LAP, CAT, KML, CPC, ARW and ZAG analyzed data. LAP, CAT, KML, CPC, ARW and ZAG interpreted results of experiments. LAP, KML and ZAG prepared figures. LAP and ZAG drafted the manuscript. LAP, CAT, KML, CPC, ARW and ZAG edited and revised manuscript. LAP, CAT, LH, KML, CPC, ARW and ZAG approved the final version of the manuscript.

## Notes

https://figshare.com/s/0cdef103fd8f43843bb4

https://figshare.com/s/3b6ae885cb2ee17b22d5

https://figshare.com/s/04384f17f5e5f603094b

https://figshare.com/s/d781f7cf7e185410c595

https://figshare.com/s/af57bff7bdf3f5dc6a45

https://figshare.com/s/06a7038abf59b1e5b9e2

https://figshare.com/s/63681359100eeb32b15a

https://figshare.com/s/a7088905168e9309adcf

https://figshare.com/s/4c1edaf591d9e999afc2

https://figshare.com/s/71fd8345296834deb3da

